# Modeling intracranial electrodes

**DOI:** 10.1101/2021.04.28.441828

**Authors:** Alejandro Blenkmann, Anne-Kristin Solbakk, Jugoslav Ivanovic, Pål Gunnar Larsson, Robert T. Knight, Tor Endestad

**Affiliations:** Department of Psychology, University of Oslo, Norway; RITMO Centre for Interdisciplinary Studies in Rhythm, Time and Motion, University of Oslo, Norway; Department of Neurosurgery, Oslo University Hospital, Norway; Department of Neuropsychology, Helgeland Hospital, Mosjøen, Norway; Department of Psychology and the Helen Wills Neuroscience Institute, University of California, Berkeley, USA

**Author notes:** **Author E-mail addresses:** Alejandro Blenkmann –. Corresponding author: Alejandro Blenkmann. Postboks 1133 Blindern 0318 OSLO.

**Keywords:** SEEG, ECoG, iEEG, intracranial electrodes, depth electrodes, subcortical grids, subdural grids, simulations

## Abstract

**Background:** Intracranial electrodes are implanted in patients with drug-resistant epilepsy as part of their pre-surgical evaluation. This allows investigation of normal and pathological brain functions with excellent spatial and temporal resolution. The spatial resolution relies on methods that precisely localize the implanted electrodes in the cerebral cortex, which is critical for drawing valid anatomical inferences about brain function.

Multiple methods have been developed to localize implanted electrodes, mainly relying on pre-implantation MRI and post-implantation CT images. However, there is no standard approach to quantify the performance of these methods systematically.

The purpose of our work is to model intracranial electrodes to simulate realistic implantation scenarios, thereby providing methods to optimize localization algorithm performance.

**Results:** We implemented novel methods to model the coordinates of implanted grids, strips, and depth electrodes, as well as the CT artifacts produced by these.

We successfully modeled a large number of realistic implantation *“scenarios”*, including different sizes, inter-electrode distances, and brain areas. In total, more than 3300 grids and strips were fitted over the brain surface, and more than 850 depth electrode arrays penetrating the cortical tissue were modeled. More than 37000 simulations of electrode array CT artifacts were performed in these *“scenarios”*, mimicking the intensity profile and orientation of real artifactual voxels. Realistic artifacts were simulated by introducing different noise levels, as well as overlapping electrodes.

**Conclusions:** We successfully developed the first platform to model implanted intracranial grids, strips, and depth electrodes and realistically simulate CT artifacts and noise.

These methods set the basis for developing more complex models, while simulations allow the performance evaluation of electrode localization techniques systematically.

The methods described in this article, and the results obtained from the simulations, are freely available via open repositories. A graphical user interface implementation is also accessible via the open-source iElectrodes toolbox.

## 1. Background

Intracranial subdural grids and depth electrodes are implanted in patients with drug-resistant epilepsy as part of their pre-surgical evaluation. Electrophysiological and neuroanatomical data are used to delineate the seizure onset zone and functional areas that will guide resective surgery (Rosenow & Luders, 2001). Intracranial electroencephalography (iEEG) recordings provide insights into human brain electrophysiology and functional mapping with an unparalleled spatial and temporal resolution, offering both clinical and research applications. Knowing the exact location of electrodes in relation to the individual cortical or subcortical anatomy is a prerequisite for a complete understanding of the electrophysiological data; leading to a precise resection of the epileptic foci and the anatomical localization of specific brain functions (Lachaux et al., 2003; Parvizi & Kastner, 2017; Stolk et al., 2018; Frausher et al., 2018).

One of the most common approaches to define electrode coordinates is to localize the artifacts they produce in post-implantation computer tomography (CT) images coregistered to pre-implantation magnetic resonance imaging (MRI) scans, a procedure that can be done manually or semi-automatically. In both cases, errors associated with the procedure or the presence of noise in the images are rarely addressed. However, it is well known that other CT artifacts than the ones of interest (e.g., connection cables or clips), adjacent electrodes, overlapping grids or strips, and noise or low image resolution make precise localization problematic (Brang et al., 2016; LaPlante et al., 2016; Narizzano et al. 2017).

Of particular interest are methods to localize high-density (HD) subdural grids and depth electrodes, which have become more frequently used in the last few years (Gupta et al., 2014; Chang, 2015). The spatial resolution of these arrays can be up to 2-3 mm and will likely be higher in the future (Martin et al., 2018). Their localization of individual electrodes from the CT artifacts is harder with currently available methods given the signal-to-noise ratio of images (see novel attempts in Branco et al., 2018a; Hamilton et al., 2017; Narizzano et al. 2017, Erhardt et al., 2020).

Although methods to localize CT artifacts and co-localize them to pre-implantation MRI are common approaches, there is no reliable gold standard to quantify their precision and robustness against noise. Here, we propose a new platform to model realistic implantation scenarios and CT artifacts, enabling the systematic quantification of localization errors in electrode localization methods.

We first introduce the main characteristics observed in CT electrode artifacts and a simple model for these. We then present the methods for fitting subdural grid and strip electrode models onto the smooth hull surface, as well as depth electrode array models targeting subcortical sites. We describe the simulation of CT noise and overlapping grids and strips. We then describe the simulation results in standardized Montreal Neurological Institute (MNI) space and single subject native anatomy. Finally, we discuss the results in the context of existing electrode localization methods, limitations of our approach, and future challenges.

The methods and results presented in this article are publicly available (except for individual patient data). Unless otherwise specified, the methods were implemented in Matlab R2019 (The MathWorks Inc., USA) and iElectrodes toolbox (Blenkmann et al., 2017) for Matlab.

## 2. Implementation

### 2.1. Characterization of CT artifacts from real cases

To characterize the intensity profile of CT artifacts in real conditions, we analyzed MRI and CT images from 8 adult patients with drug-resistant epilepsy who underwent iEEG recording as part of the pre-surgical evaluation for resective surgery. One patient was implanted with subdural grids only (310 contacts, inter-electrode distance (IED) from 4 to 10 mm, Ad-Tech Medical Instrument Corporation, USA, or PMT Corp, USA). Two patients were implanted with subdural grids and depth electrodes (153 and 314 contacts per subject, IED from 4 to 10 mm, Ad-Tech Medical Instrument Corporation, USA, or PMT Corp, USA), and five patients with depth electrodes only (between 110 and 172 contacts per subject, 763 in total, IED from 3 to 4 mm, DIXI Medical, France).

We followed a routine procedure to localize intracranial electrodes (Blenkmann et al., 2019; Stolk et al., 2018). Pre-implantation T1-weighted MRI images were processed using the FreeSurfer standard pipeline (Dale, Fischl, & Sereno, 1999), where individual brain segmentation images, pial surfaces, and cortical parcellation images (Destrieux atlas) were obtained (Destrieux, Fischl, Dale, & Halgren, 2010). Post-implantation CT images were coregistered to the pre-implantation MRI using SPM 12 software (Studholme, Hill, & Hawkes, 1999). Realigned MRI and CT images were resampled to 0.5 × 0.5 × 0.5 mm resolution. CT images were thresholded to visualize the clusters of high-intensity voxels (also known as CT artifacts). Threshold values were visually defined to identify clusters of voxels representing individual electrodes following the procedure described in Blenkmann et al. (2017). The center of each cluster of high-intensity voxels was visually identified, and the cluster was extracted and stored. For each electrode, we computed its principal axis. For grids, we computed the orthogonal direction at each electrode given its closest neighbors, and for depth electrodes, we computed the principal axis as the direction connecting the first and last electrodes of the array. We aligned the principal axes to the z-axis and aligned their corresponding centers at the coordinate system’s origin.

The Smooth Cortical Envelope (SCE) surfaces were computed from the pial surfaces following the steps provided in section 2.3.

Figure 1 shows examples of voxels’ distribution from the aligned electrodes in two grid cases (4 and 10 mm inter-electrode distance, IED) and one depth electrode array case. Observe that the voxels have an oblate and a prolate shape for depth electrodes, mimicking the metallic contacts’ disc and cylinder shapes, respectively.

**Figure 1.**
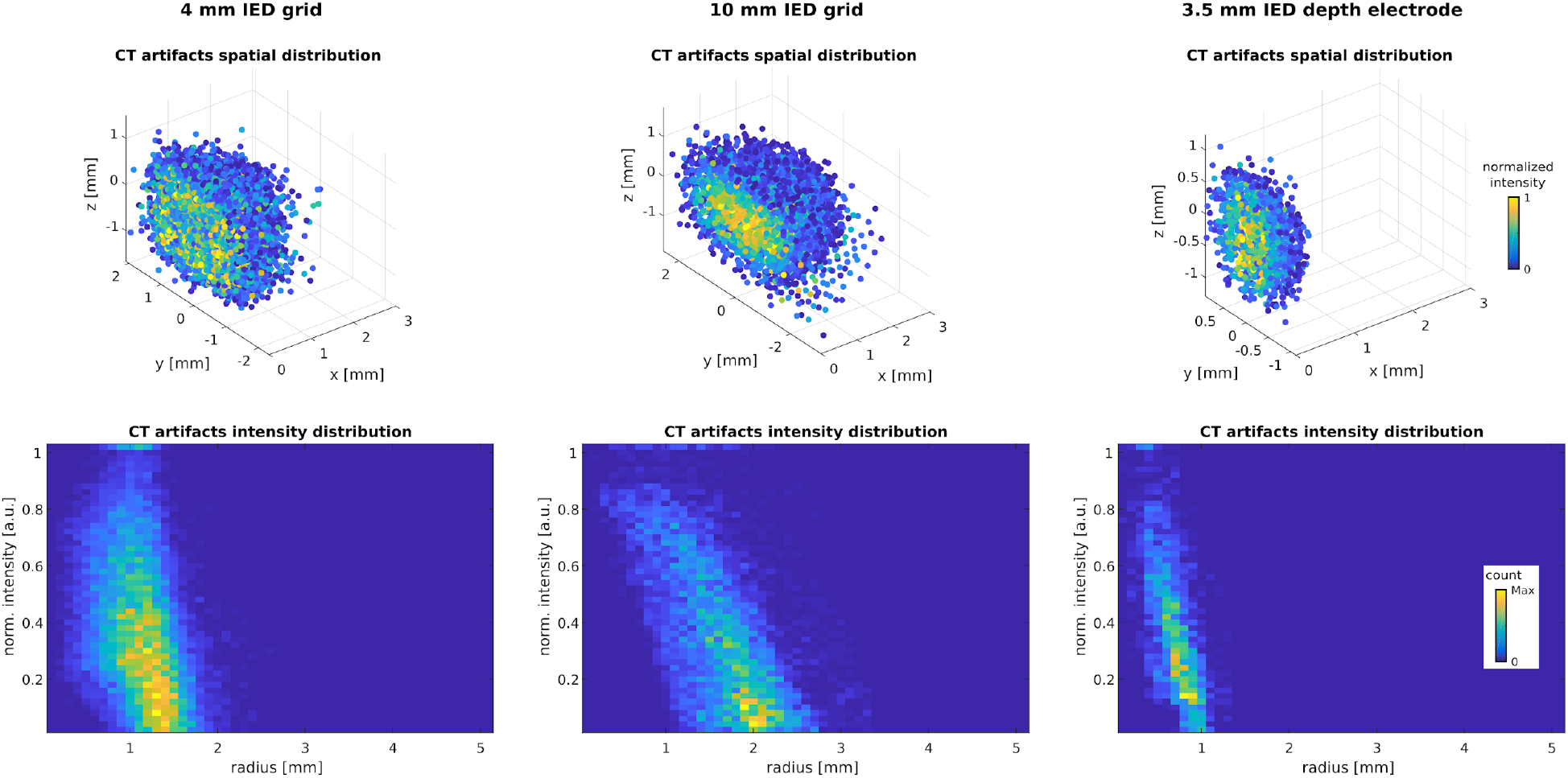
Spatial distribution of CT artifacts in three exemplary patient cases. Left column, electrodes from a subdural grid with an IED of 4 mm. Middle column, electrodes from a grid with an IED of 10 mm. Right column, depth electrodes with 3.5 mm IED. The top row shows the CT artifacts distributed in space. All individual electrodes were recentered at the origin (0,0,0). The principal axes (orthogonally to the cortical surface for grids) or main axes (connecting outer electrodes for depths) were aligned to the z-axis. The color scale denotes the normalized CT signal intensity at each voxel. For visualization purposes, only half of the electrode voxels are shown. The bottom row shows the intensity vs. radius histogram. Note that intensity decreases with radius. The color scale denotes the count of voxels in the histograms. IED: Inter-Electrode Distance

We computed 2-dimensional histograms showing the bivariate distribution of voxels in terms of intensity and radial distance to the center (Figure 1, lowest row). In all cases, we observed a tendency for a linear decrease of intensity with radius.

The spatial organization of electrodes in strips precludes the computation of orthogonal vectors. However, strip electrodes are usually of the same size as grid electrodes. Therefore, the same modeling parameters were used for both types of electrodes.

### 2.2. Modeling individual electrode CT artifacts

We used 3D ellipsoids (Figure 2A) to model the spatial distribution of individual electrodes’ CT voxels. Semi-axes length parameters (*a*, *b*, and *c*) were varied depending on electrode type and inter-electrode distance, following manufacturers’ specifications and corroborated by the CT artifacts’ observed size.

**Figure 2.**
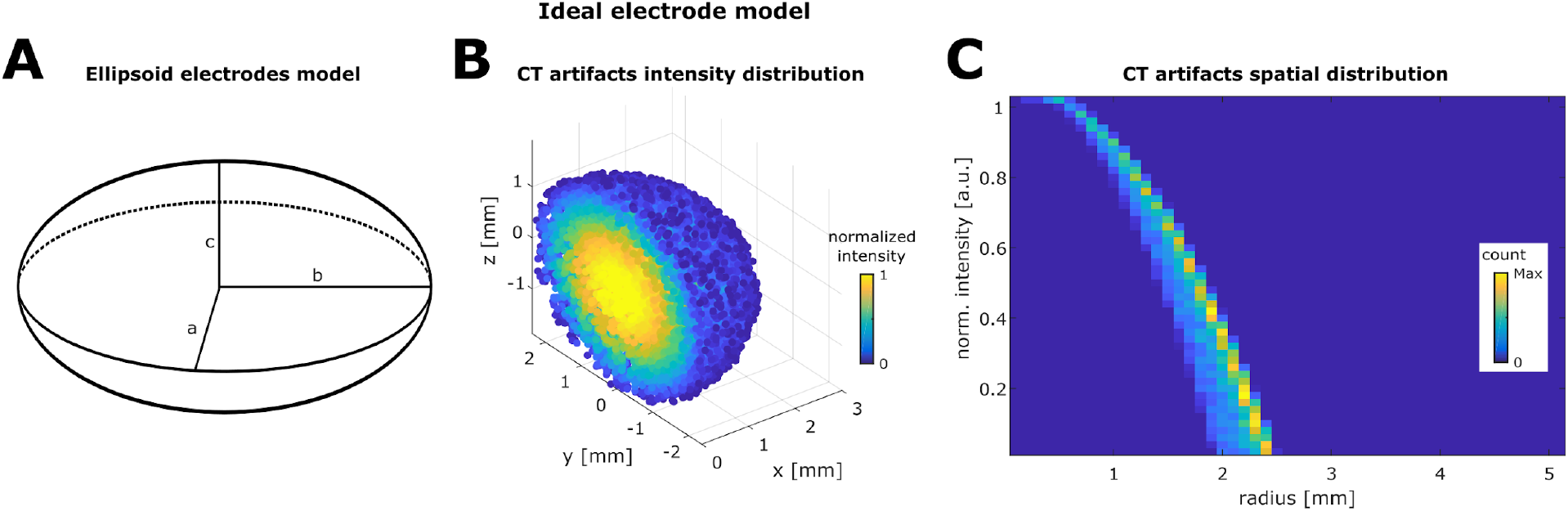
Ideal electrode model characteristics. **A.** Ellipsoid showing semi-axes model parameters.
**B.** Average distribution voxels simulated in electrodes with 10 mm IED (obtained from 64 electrodes in a grid model). The semi-axis in the z-direction is the smallest of the three, while the x and y semi-axes are the same length, producing an oblate ellipsoid shape. The intensity of each voxel is color-coded in the [0 1] range.
**C.** Intensity vs. radius histogram. The intensity profile decreases with the increasing radius, resembling the real electrode main characteristic (Figure 1). However, note that the variance observed in the ideal case histogram does not resemble the one observed in real cases.
IED: Inter-Electrode Distance

To represent the effect of discrete sampling in CT images, voxels were sampled using a 0.5 mm resolution 3D square lattice grid. To reproduce realistic CT artifacts, we used a random center location and orientation for the sampling grid. Each voxel was assigned with an intensity value depending on its location *(x, y, z)* within the ellipsoid. We defined an intensity function that declined with the radial distance as:

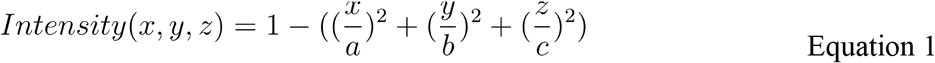

where *a*, *b*, and *c* are the semi-axes of the ellipsoid model (Figure 2A).

Figure 2B, and C shows an example of simulated grid electrodes, where the shape and intensity distribution of the voxels with the radius are similar to those observed in the real cases.

### 2.3. The Smooth Cortical Envelope surface, seed points, entry points, and target points

The implantation of real grids and strips onto different cortical areas introduces variations in how these arrays bend to follow the brain curvature. To reproduce realistic scenarios for grid implantations, we simulated grids implanted over 3D cortical surfaces. We used high-resolution pial surfaces extracted from the MNI atlas as a starting point (or individual structural MRI images for simulations in native space; see section 2.8 for details). In our processing pipeline, we used Freesurfer software (Dale et al., 1999), and individual cortical parcellation images were obtained using the Destrieux atlas (Destrieux, Fischl, Dale, & Halgren, 2010), but other atlases or software could also be used. We then computed a Smooth Cortical Envelope (SCE) surface for each cerebral hemisphere by enclosing the corresponding pial surface with a 30 mm radius sphere. A mesh smoothing was applied to remove small local protuberances (low-pass spatial filter, 100 iterations, alpha weight = 0.5, Iso2Mesh toolbox, Fang & Boas, 2009). Sup. Figure 1 shows an example of the pial surface and SCE computed for an individual subject’s brain.

We visually selected seed points over the SCE surface and used them as reference points to model grids, strips, and depth electrode arrays. We computed the local curvature of a smoothed version of the SCE (Rusinkiewicz, 2004). Seed points were visually selected on regions where the local curvature was relatively *“Low”*, *“Medium”*, or *“High”* within the range of curvature values. Depending on the location and the local curvature of the SCE surface, we modeled different electrode arrays.

In clinical practice, the implantation of depth electrodes is defined by two points, and therefore a unique trajectory connecting them. The points are typically defined as an “entry point” on the cortical surface, and a “target point” at the deepest brain location reached. We will adopt this nomenclature throughout this paper.

It is common practice to use trajectories orthogonal to the skull surface to avoid sliding of the drill and minimize bone damage during surgery. Therefore, given entry points on the SCE surface, we defined trajectory vectors orthogonal to this surface for depth electrodes simulations. Target points were defined as the deepest electrode point in the trajectory within the brain tissue (enclosed by the SCE surface). In some cases, the electrode arrays were shorter than the amount of tissue crossed by the trajectory. In these scenarios, the target was re-defined as a random point within the trajectory, while keeping all contacts inside the brain.

### 2.4. Modeling electrode array coordinates

The following steps are performed for:

- Modeling subdural grids
  1. A 2D flat model is generated, given the number of rows, the number of columns, and the inter-electrode distance parameters. We compute the center coordinate of each electrode contact.
  2. The 2D model is placed tangentially to the SCE in a given “seed” point (Figure 3D). The center of the array is aligned to the seed point.
  3. The 2D model is fitted over the SCE surface using an energy minimization algorithm (Figure 3D, Dykstra et al., 2012; Trotta et al., 2018). Briefly, an energy function:

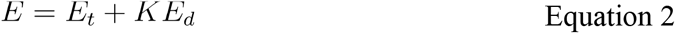

is minimized, where *E_t_* is the translation energy, *E_d_* is the deformation energy, and *K* is a constant value. The energy minimization is constrained to the electrode coordinates *x_i_* being closely located over the SCE surface:

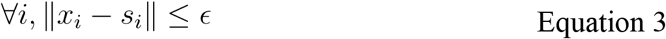

where *s_i_* is the closest point in the SCE surface to electrode *i*, and ε is a tolerance distance. *E_*t*_* (Figure 3B) represents the energy needed to translate the electrode coordinates from the original position of each electrode 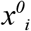 to the final position *x_i_* on the SCE surface:

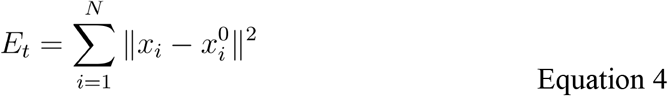

where N is the number of electrodes. *E_*d*_* (Figure 3C) represents the energy required to change the inter-electrode distance between neighbors from the original to the final location.

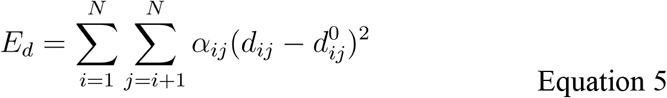

where *d_*ij*_* is the distance between electrodes, 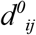 is the original distance between contacts, and *α_*ij*_* takes values of 1 or 0 if electrodes *i* and *j* are neighbors or not, respectively. To control for normal, bending, and shear deformations, first, second, and diagonal (grids only) neighbors are considered as shown in Figure 3A (Trotta et al., 2017). *E_*d*_* is typically interpreted as the deformation of “springs” connecting the electrodes (Dykstra et al., 2012).
  4. A normal vector at each electrode coordinate is computed. We select the coordinates from each electrode of interest and the nearest neighbors and compute a principal component analysis (PCA) of these points. The normal vector is one associated with the smallest component (Figure 3E).
- Modeling subdural strips
  1. A 2D flat model is generated, given the number of contacts (columns), and inter-electrode distance. The number of rows is set to three to avoid unrealistic geometrical deformations (e.g., “snake” shapes).
  2. This step is the same as step 2 for grids.
  3. This step is the same as step 3 for grids.
  4. This step is the same as step 4 for grids.
  5. Only the middle row coordinates are kept from the three rows, while the lateral ones are discarded. The result is a set of coordinates for the 1 × columns contacts.

**Figure 3.**
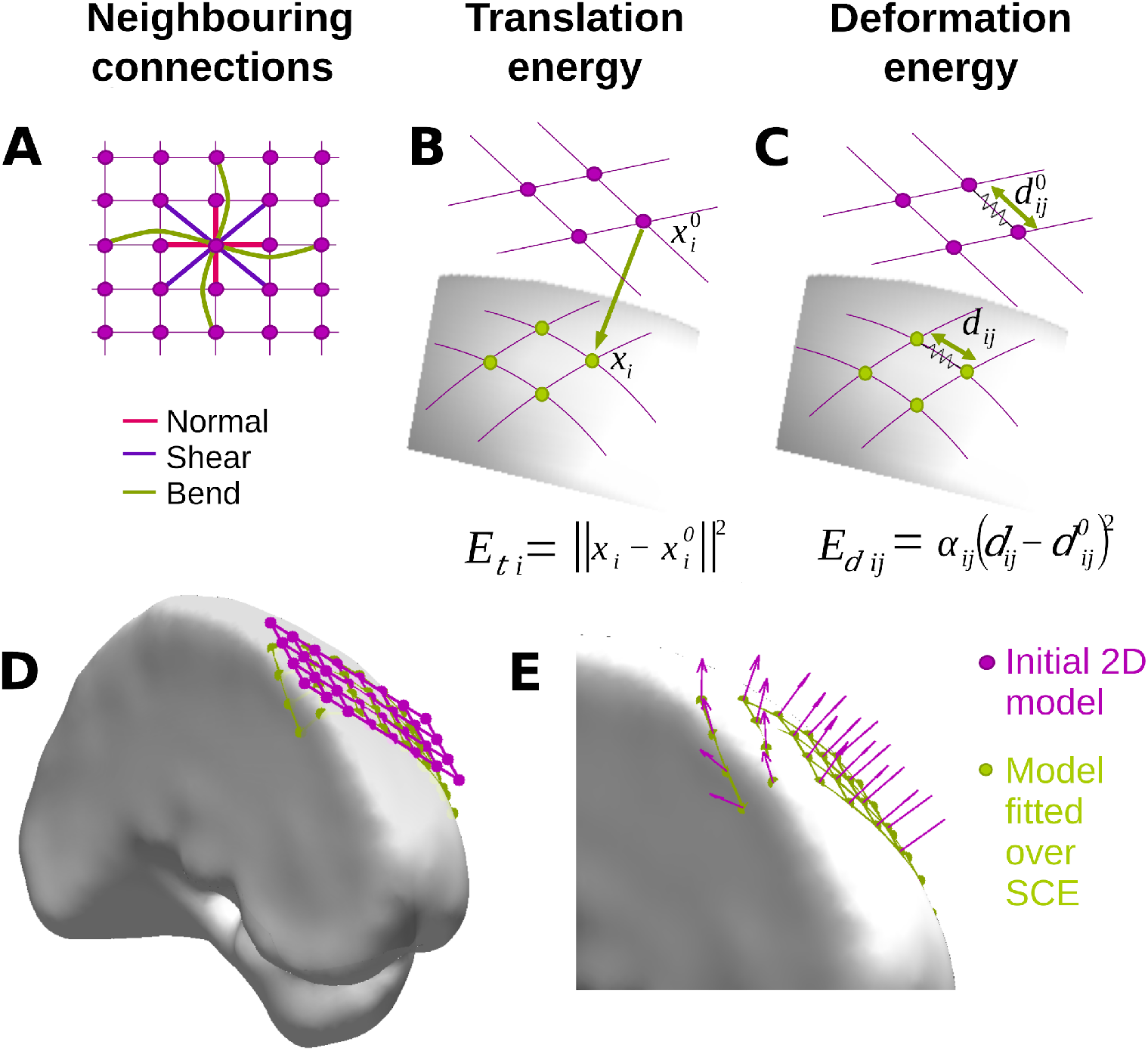
Modeling grid electrode coordinates over the SCE surface. **A.** Diagram showing the neighboring connections used to compute the deformation energy E_d._.
**B.** The translation energy E_t i_ associated with electrode i is proportional to the distance between the original 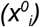 and the final location (x_i_) over the SCE surface.
**C.** The deformation energy E_d ij_ associated with neighboring electrodes i and j is proportional to the amount of deformation between the original 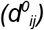 and final distance (d_ij_). Equation details are explained in section 2.4.
**D.** 2D grid model (4×8, 10 mm IED) before (pink) and after (green) being fitted to the SCE surface on the left frontal lobe.
**E.** Zoom-in detail showing the normal vectors at each electrode location. Normal vectors are used afterward to orient the simulated CT artifacts.
IED: Inter-Electrode Distance, SCE: Smooth Cortical Envelope

Deformations are needed to fit a grid or strip (originally plane objects) over the brain surface (a curved object). When localizing grids, we typically observe that these deformations are small. Therefore, we measured the IED after fitting grids and strips to the SCE surfaces and discarded simulations if one or more neighboring electrodes had IED variations over 5% from the original values.

- Modeling depth electrode arrays
  1. A unidimensional model of the electrodes is computed, uniformly distributing the contacts over the x-axis.
  2. A curvature deformation can be applied (optional) using the symmetric Lanczos window, and an arc shape electrode array can be obtained in a random orientation within a 2D plane orthogonal to the x-axis. A maximum deformation parameter η is defined as a percentage of the electrode length. η determines the maximum distance of the arc to the original model axis.
  3. The array model is aligned with the trajectory vector and translated to the electrode target point within the brain tissue.
  4. If electrode arrays are longer than the crossed brain tissue, then the deepest electrode is located on the target coordinate, contacts span along the trajectory, and contacts outside the brain tissue are discarded.

### 2.5. Modeling CT artifacts at electrode coordinates

We modeled CT artifacts for each contact in the arrays as described in section 2.2. The CT artifacts were individually computed and then aligned using the normal vector at each electrode coordinate (Figure 3E). This procedure generates CT artifacts that realistically follow the brain’s surface curvature. Figure 4A, B, and C show examples of grid and strip electrode CT artifacts over the SCE surface, whereas Figure 4D shows a depth electrode example.

**Figure 4.**
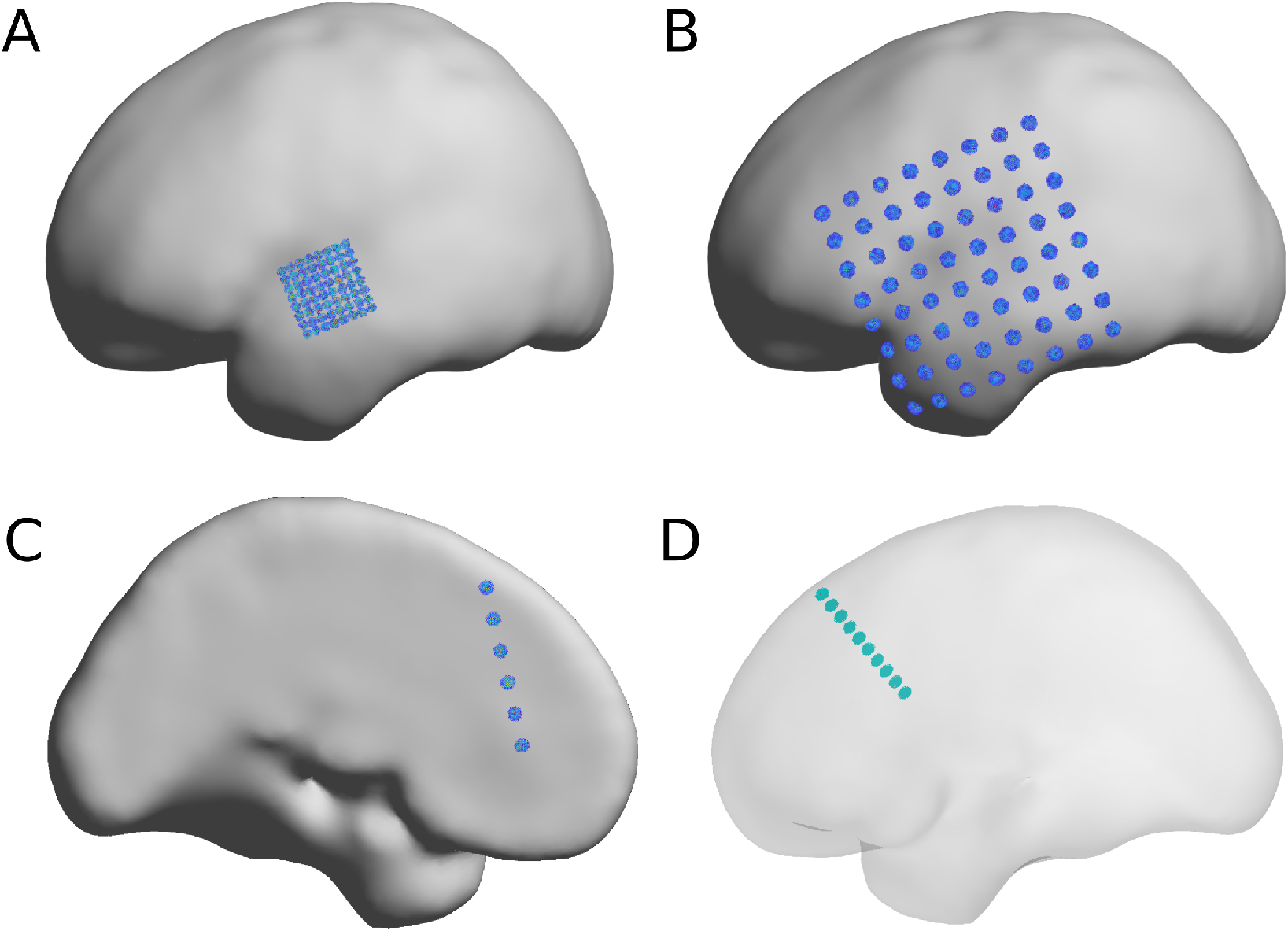
Modeled CT artifacts. Exemplary cases of modeled CT artifacts for grid and strips over the SCE, and depth electrodes penetrating the brain tissue. **A.** High-density 8×8, 3 mm IED grid over the left lateral temporal cortex.
**B.** An 8×8, 10 mm IED grid over the fronto-temporo-parietal cortex. Observe that CT voxel artifacts follow the SCE surface curvature.
**C**. Six contacts 10 mm IED strip over the left medial frontal lobe and anterior cingulate cortex.
**D**. Ten contacts, 5 mm IED depth electrodes penetrating the left frontal cortex. For illustration purposes, the SCE surface is semi-transparent.
IED: Inter-Electrode Distance, SCE: Smooth Cortical Envelope

### 2.6. Modeling CT artifacts’ noise

The localization of intracranial electrodes is typically sensitive to the signal-to-noise ratio of the processed CT images. To model the noise in the CT artifacts, we randomly displaced each voxel’s original coordinate *v_0_* to a new location *v_new_*. We computed *v_new_ = v_0_ + d_rand_ l_rand_*, where *d_rand_* is a random direction vector with unitary magnitude, *l_rand_* is a scalar value obtained from a random uniform distribution in the [*0*, *IED * σ*] interval, and *σ* is a constant parameter. In this way, changing the value of *σ* generated different noise levels in the simulated CT artifacts.

### 2.7. Modeling overlapping grids and strips

Overlapping grids or strips provide an obstacle for methods that aim to detect intracranial electrodes automatically. The following steps are performed to model overlaps:

1. A percentage of overlap is defined a priori (e.g., 10%), which sets the *number of overlapping electrodes* (NOE).
2. To define spatial rotations, a set of reference points are needed. These are the same electrode coordinate points in the case of grids or the SCE surface points in the vicinity of the electrodes (15 mm radius) in the case of strips.
3. Reference points are projected to a principal component (PC) space (Figure 5A), where the third component (PC3) has the lowest variance. In the case of strips, the same transformation is applied to the electrode coordinates.
4. A 2D surface *S_fit_* is fitted to the reference points using a local linear regression algorithm (Matlab *fit* function using ‘*lowess’* option, Figure 5A).
5. Within a 2D space defined by the first two PCs, overlapping grids or strips are defined over the original array and with a given orientation (Figure 5B).
6. The overlapping electrode array is stepwise translated:
  a. The overlapping array is translated (with a step size of IED/100) in a defined outward direction (Figure 5C).
  b. The number of overlapping electrodes in the original electrodes’ vicinity is counted in each step (within a distance equal to IED √2).
  c. The translation stops when the number of overlapping electrodes reaches the NOE (Figure 5C); otherwise, repeat all prior steps.
7. Electrodes from the overlapping array that are outside the main array area are discarded.
8. The third dimension (PC3) of the overlapping array is computed using the *S_fit_* surface.
9. A space between arrays is added to represent their thickness (1 mm in the PC3 dimension).
10. Electrode coordinates are projected back to the original 3D space (Figure 5D).
11. The normal vector at each electrode is computed as the mean of the closest normal vectors from the original array, weighted by their distance.
12. CT artifacts are modeled at each electrode coordinate and aligned to the normal vector (Figure 5E).

**Figure 5.**
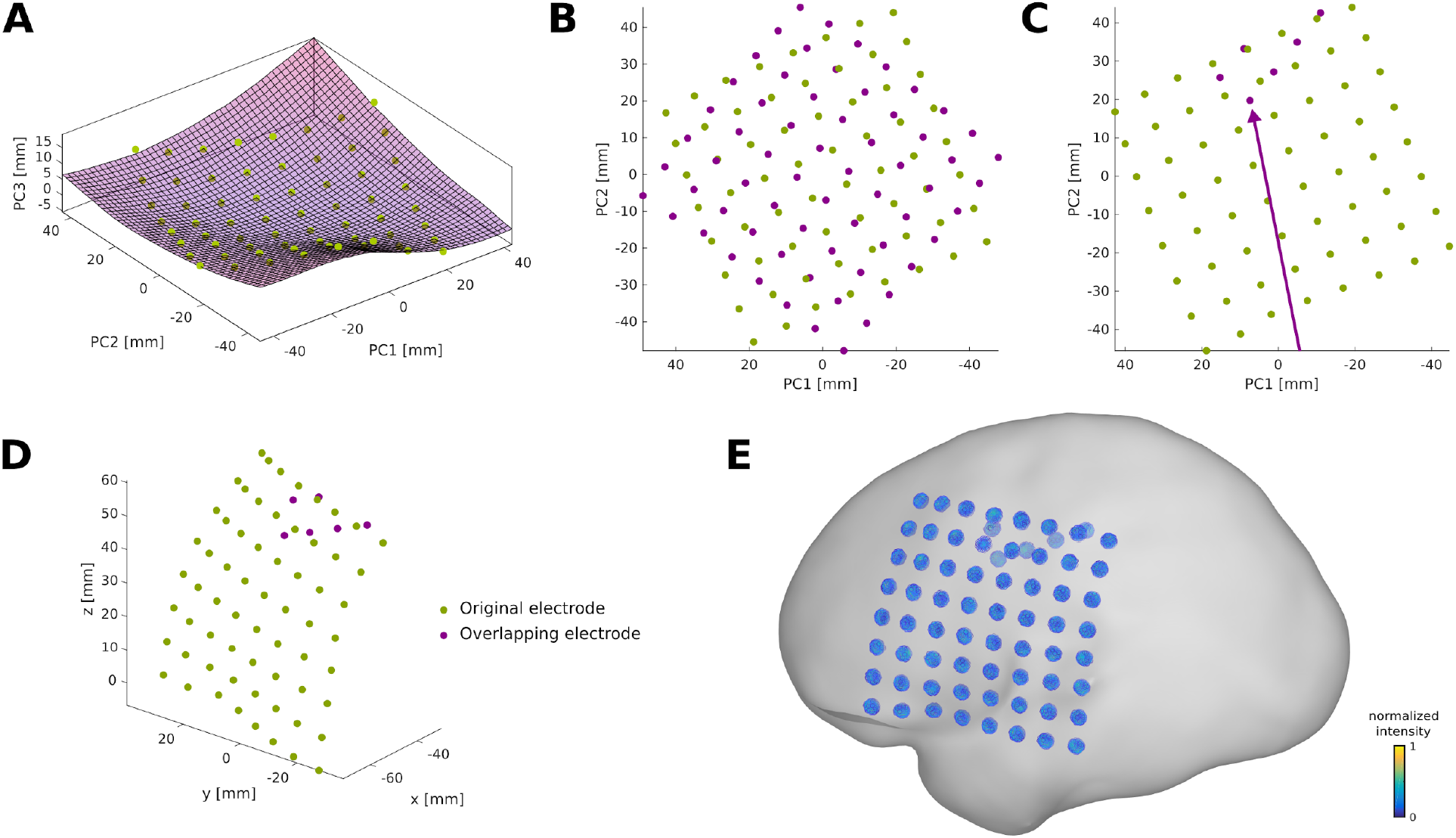
Overlapping grid. Example of an overlapping grid in an 8×8 10 mm IED case. **A.** Original electrode coordinates (green) are transformed into the principal component space, where a surface S_fit_ is fitted to the coordinates.
**B.** In a 2D space defined by PC1 and PC2, the overlapping grid (magenta) is defined with a random orientation.
**C**. The overlapping grid is translated in the arrow direction until the desired number of overlapping electrodes is achieved.
**D**. Coordinates are back-projected to the original 3D space.
**E**. Voxel artifacts are simulated in the new overlapping grid electrode coordinates (using a low noise level of σ = 0.1), overlaying the SCE surface. Voxel artifacts orientations are interpolated from close-by electrodes of the original array.
Note: for illustrative purposes, plots B and C show the PC1 axes direction inverted.
IED: Inter-Electrode Distance, SCE: Smooth Cortical Envelope

### 2.8. Modeling implanted electrodes in real subject’s space

To demonstrate the usefulness of the algorithm, we applied our methods in real native brain space. We processed pre-implantation MRI and post-implantation CT images from an adult patient with drug-resistant epilepsy, a potential candidate for resective surgery, as described in section 2.1. The SCE surfaces for each hemisphere were computed following the steps described in section 2.3.

Grids, strips, and depth electrode array coordinates were modeled using the graphical interface provided by the iElectrodes toolbox (Blenkmann et al., 2017). For grids and strips, the 2D model arrays are first manually translated and rotated over the SCE surface until the desired location is reached and fitted to the SCE surface. Depth electrodes are defined by manually setting the target and entry points.

## 3. Results

### 3.1. Grids and strips

We simulated electrode arrays of multiple dimensions representing models frequently available in the market by different manufacturers. These cover various combinations of size and IED, including grids of 2×4, 4×4, 4×8, 8×8, 8×16, and 16×16 contacts, strips of 1×4, 1×6, and 1×8 contacts. We selected 57 seed points in the standardized MNI SCE surface with different local curvatures (*Low*, *Medium*, and *High*; Sup. Figure 2).

Grids and strips implantation scenarios were simulated several times at each seed point, rotating the grids in angles multiple of 30 degrees around the center. The location and the local curvature of the SCE surface surrounding the seed points determined which arrays were simulated. Large grids were simulated on *Low* curvature regions, whereas medium and small size grids were simulated over *Medium* and *High* curvature areas. For example, over the lateral fronto-temporo-parietal cortex, it is realistic to simulate an 8×8, 10 mm IED grid, but unrealistic to simulate the implantation of such a big grid over the frontal pole.

For the fitting of grid and strips onto the SCE surface, coefficient *K* was set to 1000, and the tolerance distance ε = 0.1 mm for all arrays, except for the 16×16 grid cases where *K =* 100 and ε = 0.5 mm (Eq. 2).

A total of 3646 scenarios for grids and strips were simulated and in 3321 instances the arrays were successfully placed over the SCE surface. In 9% of the cases, the IED deformations between at least one pair of contacts exceeded the 5% threshold and were discarded.

Finally, we simulated the CT artifacts at each electrode location and manipulated the noise levels with *σ =* 0, 0.1, 0.2, 0.3, and 0.4. Moreover, grids and strips were simulated with a 10% overlap and no overlap. Overlapping grids were defined with a random orientation and translated in a random direction. In principle, grids and strips of any size and IED can be overlapped. However, for the sake of simplicity, we used arrays with the same dimensions.

Altogether, we produced approximately 33000 simulations of CT artifacts. Figure 7 shows how high- and low-noise levels affect the voxels’ distribution in space for high- and low-density grids (left and middle columns, respectively). Notice that *σ =* 0.1 (low-noise) generates an intensity histogram that resembles a real low-noise scenario (Figure 2).

To evaluate the simulated coordinates’ quality, we quantified the deformations introduced by the grids’ projection onto the SCE. We measured the distance between 1^st^, 2^nd^, and diagonal neighboring contacts, normalized in each case by the IED. Additionally, we measured the distance between the fitted electrodes and the SCE.

Overall, the median deformation for 1^st^, 2^nd^, and diagonal neighbors were 0.174, 0.169, and 0.243 % of the IED. The median distance of electrodes to the SCE was 0.012 mm.

Figure 6A shows that median deformation increases for higher values of IED and SCE curvature. Figure 6B shows that the electrodes’ median distance to the SCE decreases with the number of contacts and increases with the SCE curvature.

**Figure 6.**
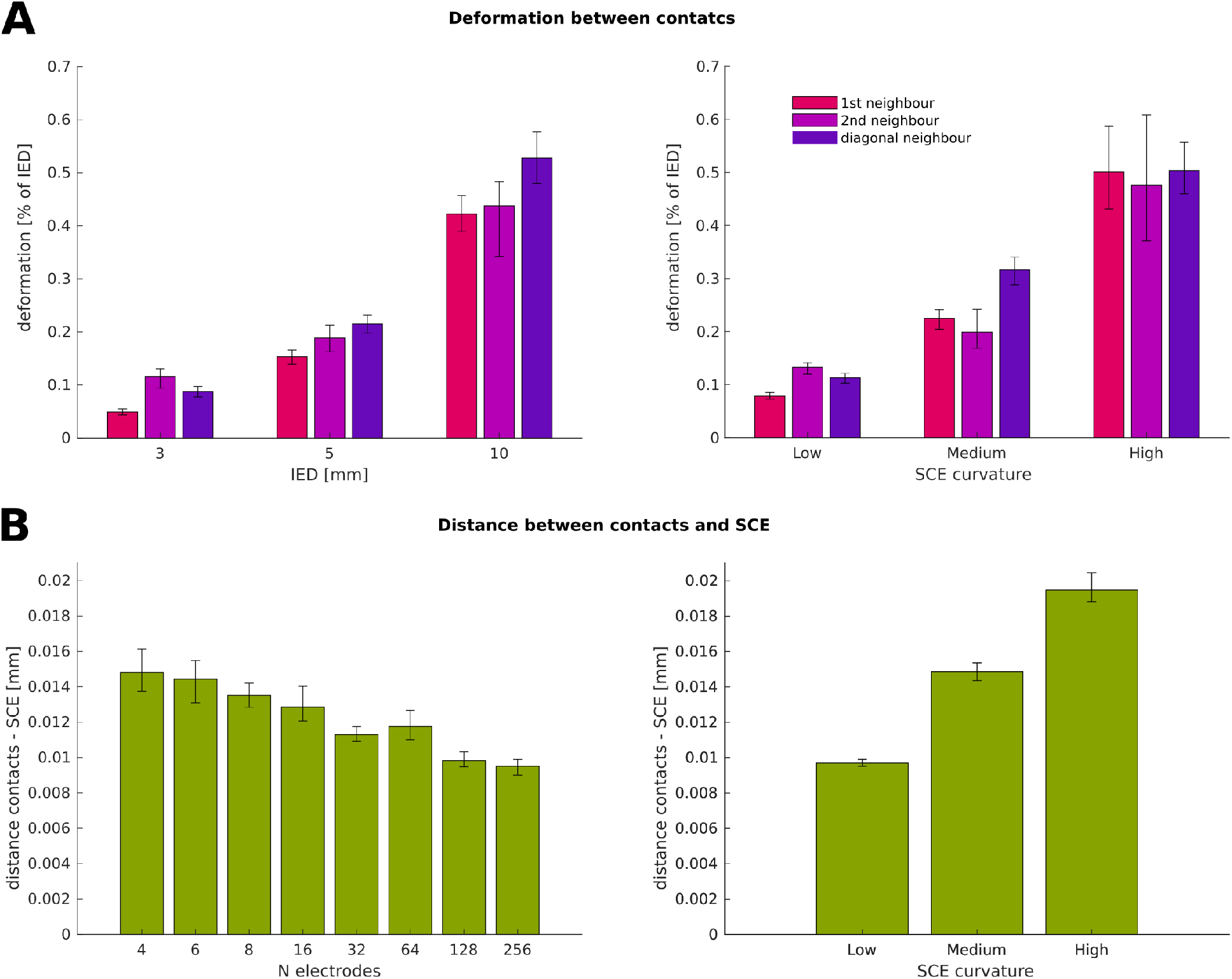
Deformation characteristics of the simulated grids and strips. **A**: Median deformation between contacts as a function of IED (left) and SCE curvature (right).
An effect of IED and SCE curvature can be observed over the distance between contacts.
**B**: Median distance between electrodes and SCE as a function of the number of electrodes (left) and SCE curvature (right).
Error bars denote 95% CI of the median obtained by bootstrapping. SCE: Smooth Cortical Envelope. IED: Inter-Electrode Distance.

**Figure 7.**
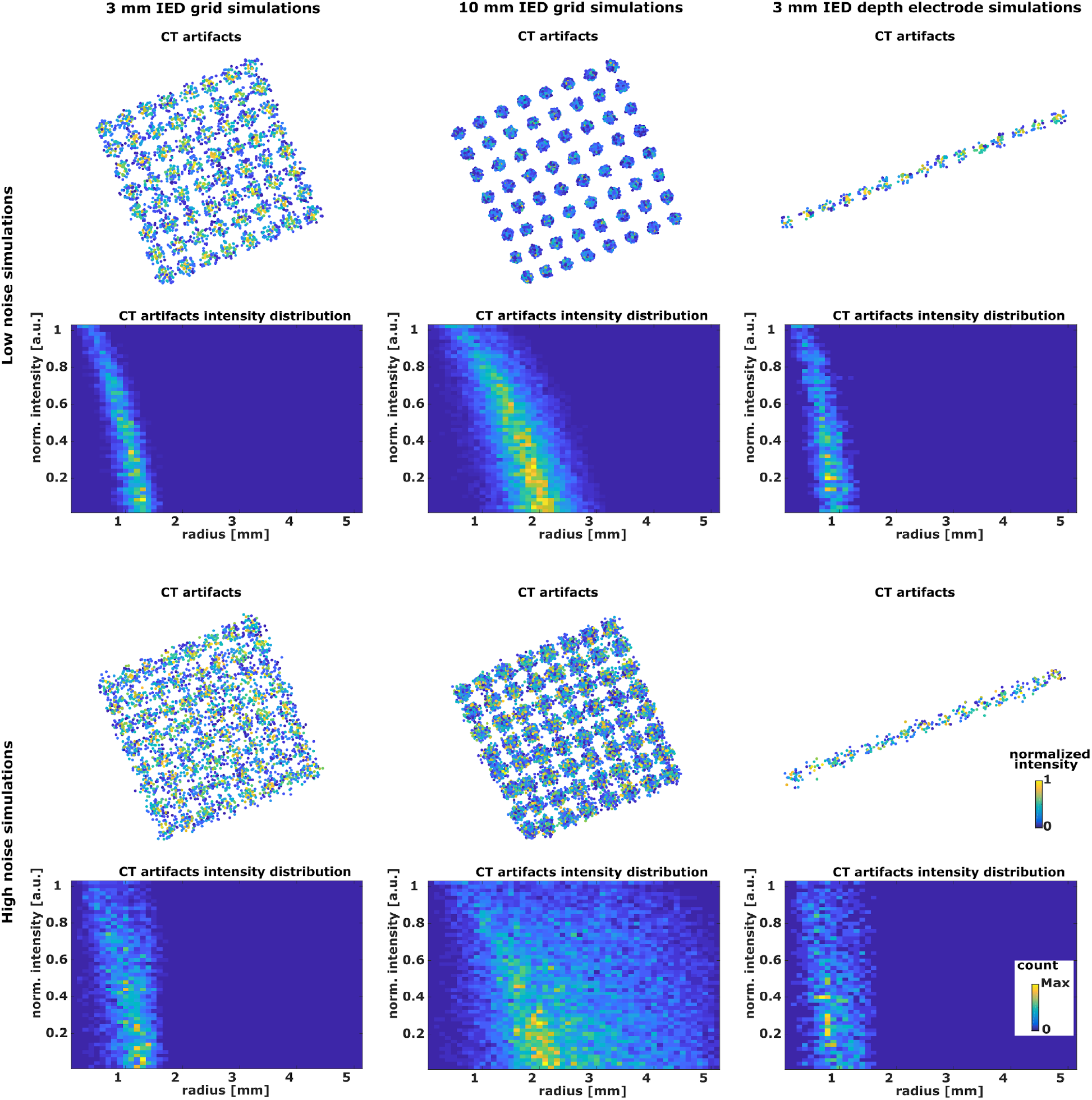
Noise effect on CT artifacts. Examples of the effect of the noise level in simulated grid and depth electrodes artifacts. Columns: First, an 8×8 3 mm IED grid; second, an 8×8 10 mm IED grid; third, a 15 contact 3 mm IED depth electrode array.
Rows: The first two rows show the results for low noise simulations (σ = 0.1), whereas the last two rows show the results for high noise simulations (σ = 0.4).
For each case, the CT artifacts are plotted on top, and intensity vs. radius histograms are shown below. Note that the spatial scale in the CT artifact plots is different for each column. IED: Inter-Electrode Distance.

### 3.2. Depth electrodes

For depth electrodes, a total of 33 entry points over the MNI SCE were defined from the previous set of 57 seed points, restricted to locations where the implantation of depth electrodes is realistic. At each point, we simulated the implantation of depth electrodes as combinations of 4, 8, 10, 12, 15, and 18 contacts; 3, 5, and 10 mm of IED; and linear or curved deformation (maximum deformation η of 1% of the total length). This resulted in a total of 858 simulation scenarios for depth electrodes within the MNI brain. Then, we simulated CT artifacts for each scenario, manipulating the noise levels in the same way we did for grids and strips. We obtained a total of 4290 CT artifact array simulations. Figure 7 (right column) shows how low- and high-noise levels affect the voxels’ distribution in space.

### 3.3. Simulations in real subject’s anatomical space

To assess the usefulness of our platform, we tested our algorithms on a real subject’s anatomical brain space. We simulated grids, strips, and depth electrodes over multiple center points (seeds) and orientations. After the array center’s initial definition, dedicated controls in the graphical interface permit accurate center and orientation changes. This allows the user to precisely locate the electrodes in the desired position.

Figure 8 shows an example of multiple electrode arrays simulated over the pial surface in a single subject. A lateral high-density grid (5 mm IED) was simulated over the right fronto-temporo-parietal region, and its fifth row was particularly aligned with the superior temporal gyrus. Smaller grids were simulated over the fronto-parietal cortex. Strips were simulated to cover the anterior lateral frontal and orbitofrontal cortex, and the middle and inferior temporal gyrus. Note that grid electrodes were not forced to contact the pial- but the SCE surface.

**Figure 8.**
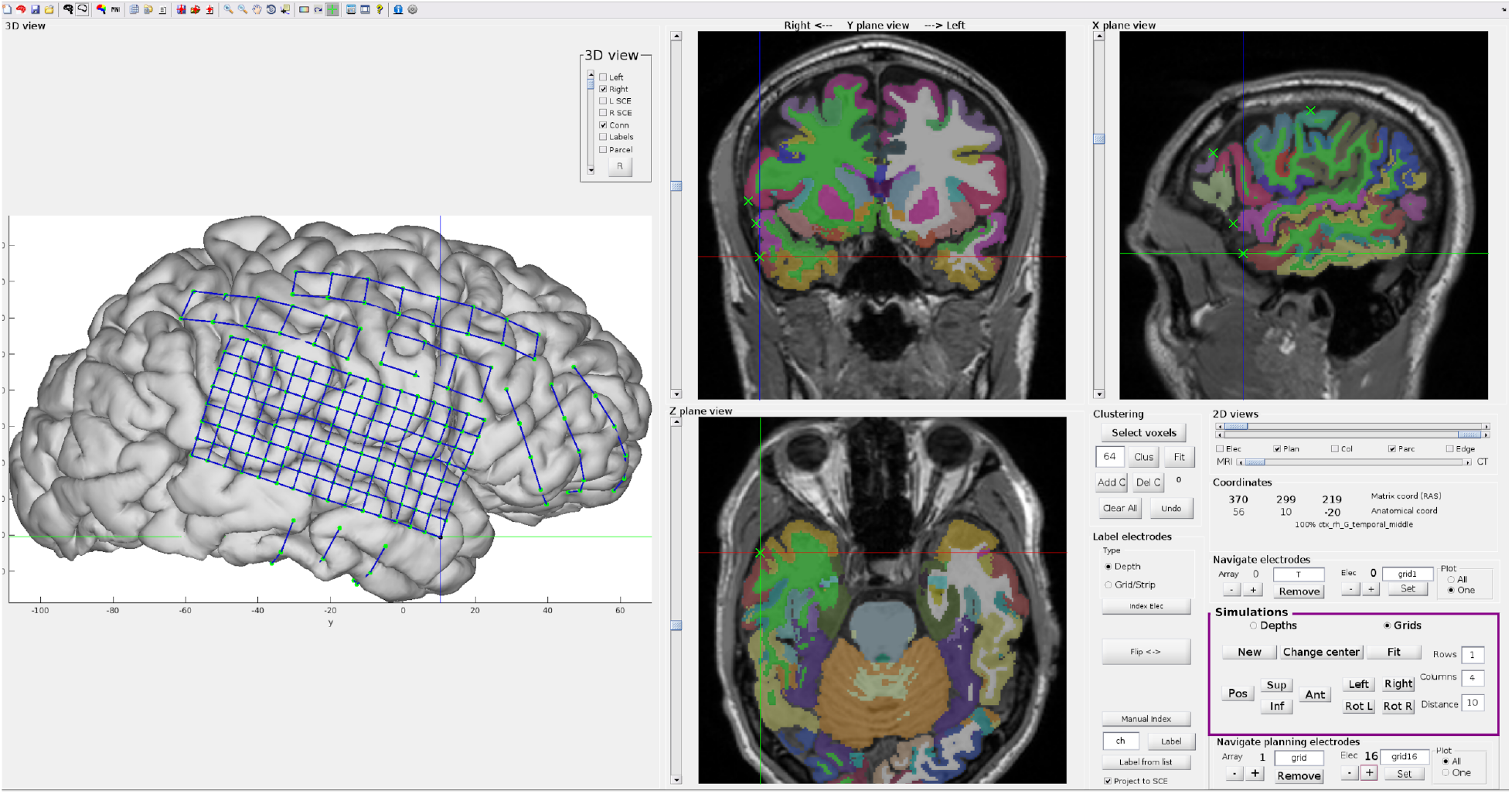
Intracranial electrode simulations on a real subject’s brain anatomy. Example of electrodes simulated on a real (native space) brain anatomy using iElectrodes graphical interface. Grids and strips were simulated covering the right cerebral hemisphere.
2D slices show the Destrieux atlas on top of the T1-weighted MRI image and simulated coordinates as green ‘x’. The box (in magenta) delineates the controls used for displacing, rotating 2D grid models, and fitting them to the SCE surface for simulation purposes.
Note that electrodes were fitted to the SCE surface, but the pial brain surface is shown instead for illustrative purposes. SCE: Smooth Cortical Envelope.

## 4. Discussion

With the aim of establishing a testbench for intracranial electrode localization algorithms, we provide the first modeling platform of intracranial electrodes. Overall, our simulations cover a wide range of realistic scenarios that can be useful for testing localization algorithms. An extensive database with cases is freely available for this purpose (see availability in section 6). We focused on providing control over those situations where localization algorithms might fail or encounter difficulties, such as high-density arrays (grids and depth electrodes), high noise levels, overlapping grids, or highly curved implants. Moreover, specific cases can be modeled using the graphical interface of iElectrodes or the available code files.

To achieve our aim, we developed novel methods to model intracranial EEG electrode coordinates and the CT artifacts typically produced by these. Simulated coordinates for grids and strips were obtained by fitting models to the smooth cortical envelope (SCE) surface. We simulated coordinates in cortical areas with different curvatures using arrays of various geometrical dimensions to mimic realistic scenarios. Additionally, depth electrodes were modeled between anatomical target and entry points, and precise deformations were parametrically defined for each case.

Finally, we simulated the CT artifacts at the electrodes’ coordinates. The distribution of intensity over space, the shape of the artifacts, and the orientation of each cluster of high-intensity voxels were carefully modeled. Moreover, different noise levels and overlapping electrodes were simulated, mimicking real scenarios.

### 4.1. Modeling implantation coordinates

To evaluate the accuracy of grid and strip implantation models we measured their deformation (i.e., changes in the inter-electrode distance). Fitting grids or strips over the SCE surface introduce deformations since plane- or line-shaped models have to be bent to fit the curved brain envelope surface. Importantly, deformation values were relatively small, with a median value of less than 0.3% of the IED, an indication that the resulting models are reliable.

We discarded simulations where the deformations exceeded 5% of the IED, since these cases do not represent realistic scenarios. Similar results could be achieved by constraining the solutions of the energy function minimization (Eq. 2), with the additional cost of increased computational time.

As shown in Figure 6A, grid deformations increased with larger IED. We theorize that bigger grids had to deform more to cover a larger extent of the curved SCE surface, in contrast to smaller grids that suffer less deformation in the reduced area they overlay. Similarly, higher levels of curvature produced larger deformation values (Figure 6A).

Parameter *K* (Eq. 2) controls the deformation introduced in the fitting procedure. We defined *K = 1000* which allowed very small deformations (Trotta et. al; 2018). However, for large high-density grids (IED of 3 mm) we had to reduce the value to *K = 100* since the algorithm did not reach feasible solutions in most scenarios. We suggest using the highest *K* value that reaches a feasible solution, therefore introducing the smallest deformations.

Another way to quantify the accuracy of grids and strips models is to measure the distance between electrode coordinates and the SCE, since electrodes are expected to be as close as possible to the SCE. Grid and strip electrodes fitting to the SCE were constrained by ε, a tolerance distance between these two (Eq. 3). This parameter enables direct control of the simulation quality. Overall, the resultant distances between electrodes and SCE (median of 0.012 mm) were negligible when compared to the IEDs (in the order of several mm) indicating successful algorithm performance.

Interestingly, the number of electrodes and the local curvature of the SCE surface affected this distance (Figure 6B). A higher number of electrodes allowed the distance to be smaller by following the SCE curvature more closely. The larger local curvature also introduced a larger distance between the contacts and the SCE surface. Still, the values are relatively small when compared to the arrays’ dimensions.

Depth electrode arrays, on the other hand, bend during implantation mainly due to brain shift. Our simulations successfully resulted in deformation values equivalent to the ones reported in clinical practice (~ 1-3 mm at the tip; Vakharia et al., 2017; Cardinale et al., 2013).

Our approach provides precise control of the deformation by applying an arc-shaped function that departures from the original straight linear model. In our methods, we used a symmetric Lanczos window function. However, the use of other functions is straightforward.

### 4.2. Simulation of CT artifacts, noise, and overlapping electrodes

Previously, models of individual electrode CT artifacts were done as uniform intensity cylinders (Brang et al., 2016), without considering the details examined in the current study. We used the bivariate intensity-radius histograms from real CT artifacts (Figure 1) as a guide to model realistic CT artifacts. We defined ellipsoid-shaped artifacts, with the intensity changing as a function of the radius. The noise-free ideal electrode model produced a histogram profile that resembles the real artifacts’s main characteristic, i.e., an intensity decrease with increasing radius (Figure 2). However, the histogram profile is “cleaner”, i.e., it lacks the variance or jitter observed in real cases. For this reason, the introduction of noise plays a significant role in resembling realistic histograms (Figure 7). We suggest using at least a low noise level (*σ =* 0.1) to achieve realistic simulations.

Electrode CT artifacts corresponding to grids and strips were placed over the SCE surface, keeping electrode principal axes orthogonal to the surface. Meanwhile, the ones corresponding to depth electrode arrays penetrating the brain were aligned to the arrays’ main axes. Finally, we modeled overlapping CT artifacts for grids and strips, controlling the size, orientation, and the number of overlapping electrodes. The last two features add a level of realism to the models not shown before.

### 4.3. The relevance of intracranial electrode models for localization algorithms

Over the last decade, several approaches have been proposed to localize intracranial electrodes based on CT and MRI images. The majority of approaches focused on post-implantation CT and pre-implantation MRI images. The detection of CT artifacts has typically been a manual process, but has recently been approached by semiautomatic techniques such as clustering voxels of high intensity (Blenkmann et al., 2017; Brang et al., 2016; Branco et al., 2018a; Taimouri et al., 2013; Quin et al., 2017), or the interpolation of coordinates given entry and target points in depth electrodes (Li et al., 2020; Arnulfo et al., 2015). Noise and overlapping electrodes are two well-known difficulties for these algorithms, precluding the success of automatic methods. For example, Brang and colleagues (2016) excluded overlapping electrodes from the analysis given the resulting difficulties, while others treated such cases manually (Branco et al., 2018a; LaPlante et al., 2016; Taimouri et al., 2014). In the same vein, Narizzano and colleagues (2017) observed errors in their depth electrode estimations associated with other electrodes in the proximity, requiring manual intervention from the user. Moreover, noise signals could be mistakenly detected as electrodes (La Plante et al., 2016), whereas CT image resolution affects localization accuracy (Brang et al., 2016). Apart from the studies above, the effect of signal-to-noise ratio on the precision of electrode localization algorithms was rarely discussed, most likely due to the lack of standardized measures to quantify the noise level. The proposed framework provides a controlled simulation of noise levels and overlapping electrodes, allowing performance evaluation of different localization algorithms.

The spatial resolution of grids and depth electrodes has increased over the last years (Chang, 2015; Erhardt et al., 2020), and high-density arrays are more informative than low-density ones in both cognitive (Gupta et al., 2014; Jiang et al., 2018) and clinical research (Stead et al., 2010). High-density electrodes require additional spatial precision and can be an obstacle for many of the frequently used localization algorithms (but see: Branco et al., 2018b; Hamilton et al., 2017; Narizzano et al., 2017). Simulations can be a reliable platform to develop novel localization techniques for high-density electrode arrays (e.g., Blenkmann et al., unpublished).

It is important to mention that our models introduced deformations on the order of 1/1000 of the inter-electrode distance, and distances between electrodes and the SCE on the order of 1/100 mm, which ensures precise modeling of the grids and strips. These errors and deformations are negligible compared with those observed with previous localization (~ 0.2-0.6 mm; Blenkmann et al., 2017; Narizzano et al., 2017) and brain-shift correction algorithms (~ 2-3 mm; Trotta et al., 2018; Branco et al., 2018a; Branco et al., 2018b; Brang et al., 2016).

### 4.4. Assumptions, limitations, and future directions

Although we provide a substantial number of scenarios and a good starting point to model implanted electrodes, there are some noteworthy limitations of the current models.

First, we took a simple approach to the spatial intensity distribution of CT artifacts. The simplistic assumption allowed us to build realistic models of large arrays of electrodes. However, more sophisticated approaches could be implemented, considering the x-ray interaction with metallic electrodes and the effects produced in the image reconstructions (Boas & Fleishmann, 2011; Katsura et al., 2018). Developments in this direction could pave the way to model the artifacts produced by microwires at the tip of depth electrodes (e.g., Behnke-Fried electrodes, Ad-Tech Medical) and the design of novel localization algorithms for this specific and unsolved problem.

Second, the implantation of intracranial grids and strips is a procedure that results in the deformation of the brain tissue. Deformations of 10 mm or more can occur on the brain surface around the electrodes or in deeper brain structures due to cerebrospinal fluid loss in the ventricles (Studholme et al., 2001; LaViolette et al., 2011). Implantation of depth electrodes might also produce brain deformations but to a much lower extent, with a smaller amount of cerebrospinal fluid loss, if any (Elias et al., 2007). Modeling brain deformations is a complex problem, where multiple variables have to be considered, such as the size and location of the skull opening, the amount of cerebrospinal fluid loss, and the swelling of soft tissue, among others (Studholme et al., 2001). Given the complexity of such a problem, we assumed non-deformed brains in our simulations, precluding their use to evaluate brain-shift correction algorithms. The use of non-linear finite element methods can be a successful way to model these more complex brain deformations (Wittek et al., 2007).

## 5. Conclusions

Intracranial EEG recordings allow us to study brain function with excellent spatial resolution and rely on precisely localizing the implanted electrodes. Here, we presented the first platform to model electrode coordinates and CT artifacts of implanted grids, strips, and depth electrodes. Implanted electrodes under realistic scenarios were successfully modeled with high accuracy, resembling real cases. These methods enable the systematic and quantitative evaluation of electrode localization strategies, contributing to the development of future techniques. The platform should be considered a starting point for more sophisticated models, e.g., including brain tissue deformations or microwires.

The modeling methods and the results from the simulations are freely available to the research community via open repositories. Moreover, a graphical user interface implementation is also available via the open-source iElectrodes toolbox.

## 6. Declarations

### Ethics approval and consent to participate

Patients were recruited from the University of California, Irvine, University of California, San Francisco, and Oslo University Hospital.

This study was approved by the Regional Committees for Medical and Health Research Ethics, Region North Norway (REK 2015/175) and the Human Subjects Committees at UCSF, UC Irvine and UC Berkeley.

All patients provided written informed consent as part of the research protocol approved by the Institutional Review Board at each hospital.

### Consent for publication

Not applicable.

### Availability of data and materials

- The scripts and simulated datasets generated and analyzed during the current study are available in the OSF repository “Modeling intracranial electrodes” https://osf.io/9fsm3/ (DOI 10.17605/OSF.IO/9FSM3)
- A graphical interface for the interactive simulation of electrode coordinates is available within the iElectrodes toolbox at https://sourceforge.net/projects/ielectrodes/
- The patients’ datasets analyzed during the current study are not publicly available due to our ethical approval conditions that do not permit public archiving of anonymized study data.

### Availability and requirements

Project name: iElectrodes

Project home page: https://sourceforge.net/projects/ielectrodes/

Operating system: Platform independent

Programming language: Matlab

Other requirements: Matlab 2018a or higher (including Optimization Toolbox), iElectrodes toolbox

License: GNU GPL

### Authors’ contributions

AOB, AKS, and TE designed this study. AOB coded, and tested the software tools. AOB performed the analyses. TE, AKS, PGL, JI, and RTK provided the data and validated the results. AOB wrote the manuscript. All authors revised the manuscript. All authors read and approved the final manuscript.

## List of abbreviations

IED: Inter-Electrode Distance
SCE: Smooth Cortical Envelope

## Acknowledgments

We are grateful to the patients for kindly participating in our study. We thank FRONT Neurolab/ RITMO members for rich discussions.

## Supplementary material

**Supplementary Figure 1.**
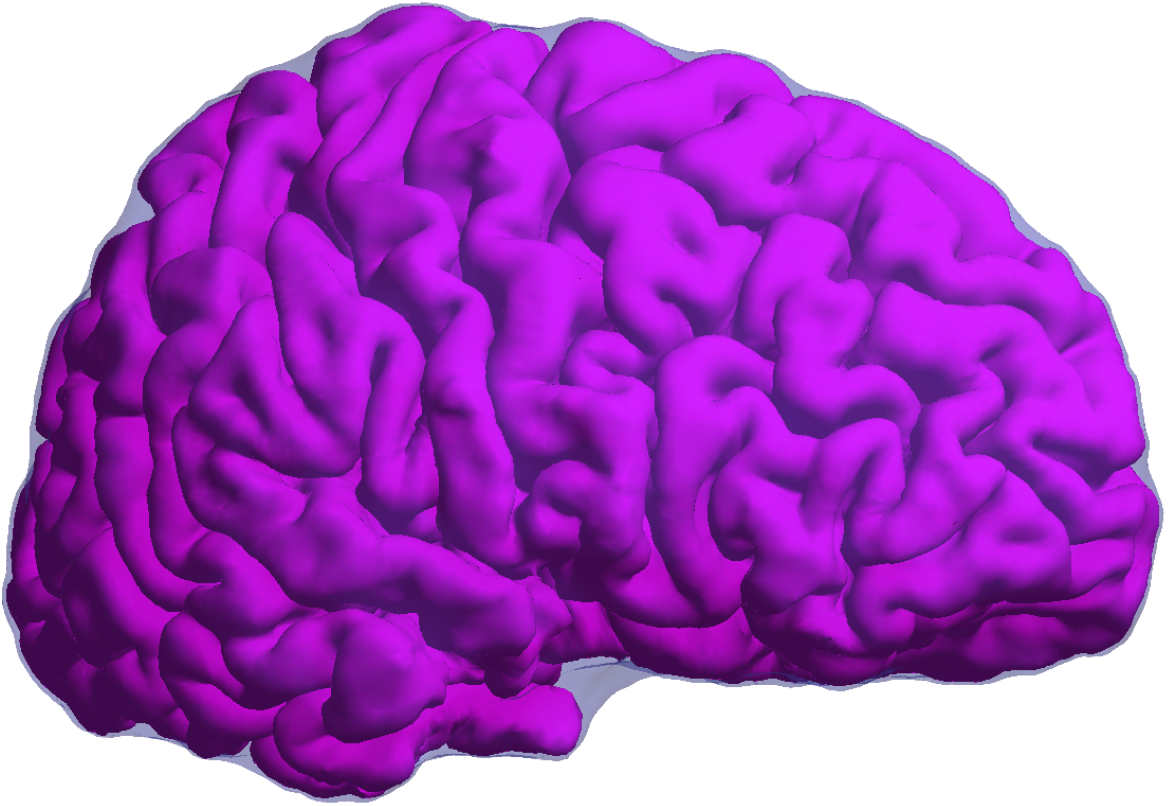
Pial surface and Smooth Cortical Envelope. Smooth Cortical Envelope (SCE, in semi-transparent violet) over the pial surface (in pink) computed for an individual subject’s brain. The SCE is enclosing the pial surface using a 30 cm sphere. Simulated grids and strips were overlaid on the SCE surface.
The same procedure was also applied to the MNI standard brain.
SCE: Smooth Cortical Envelope

**Supplementary Figure 2.**
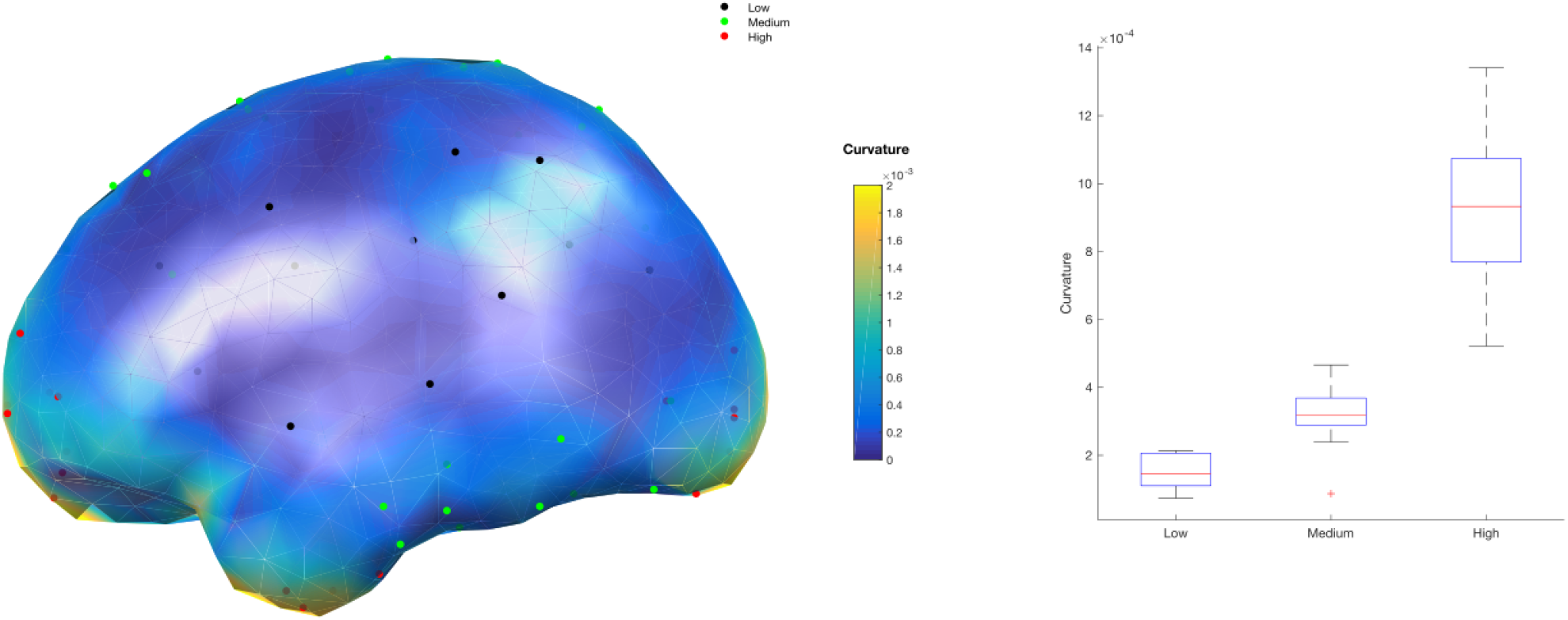
Curvature of SCE and seed points. Left: Local curvature of a smoothed version of the MNI SCE showing the selected seed points in Low (black), Medium (green), and High (red) curvature regions.
Right: Boxplots showing the curvature values for seed points in Low, Medium, and High curvature areas. Curvature values were computed for each seed point as the mean within 25 mm.
The center line (red) of each boxplot represents the median and the edges are the 25th (Q1) and 75th (Q3) percentiles. The whiskers extend to the most extreme data points not considered outliers, and the outliers are plotted individually.
SCE: Smooth Cortical Envelope

## References

Arnulfo, G., Narizzano, M., Cardinale, F., Fato, M. M., & Palva, J. M. (2015). Automatic segmentation of deep intracerebral electrodes in computed tomography scans. BMC Bioinformatics, 16(1), 1–12. https://doi.org/10.1186/s12859-015-0511-6

Blenkmann, Alejandro O., Collavini, S., Lubell, J., Llorens, A., Funderud, I., Ivanovic, J., Larsson, P. G., Meling, T. R., Bekinschtein, T., Kochen, S., Endestad, T., Knight, R. T., & Solbakk, A.-K. (2019). Auditory deviance detection in the human insula: An intracranial EEG study. Cortex, 121, 189–200. https://doi.org/10.1016/j.cortex.2019.09.002

Blenkmann, Alejandro Omar, Phillips, H. N., Princich, J. P., Rowe, J. B., Bekinschtein, T. A., Muravchik, C. H., & Kochen, S. (2017). iElectrodes: A Comprehensive Open-Source Toolbox for Depth and Subdural Grid Electrode Localization. Frontiers in Neuroinformatics, 11(March), 14. https://doi.org/10.3389/fninf.2017.00014

Boas, F. E., & Fleischmann, D. (2011). Evaluation of two iterative techniques for reducing metal artifacts in computed tomography. Radiology, 259(3), 894–902. https://doi.org/10.1148/radiol.11101782

Branco, M. P., Gaglianese, A., Glen, D. R., Hermes, D., Saad, Z. S., Petridou, N., & Ramsey, N. F. (2018a). ALICE: A tool for automatic localization of intra-cranial electrodes for clinical and high-density grids. Journal of Neuroscience Methods, 301, 43–51. https://doi.org/10.1016/j.jneumeth.2017.10.022

Branco, M. P., Leibbrand, M., Vansteensel, M. J., Freudenburg, Z. V, & Ramsey, N. F. (2018b). GridLoc: An automatic and unsupervised localization method for high-density ECoG grids. NeuroImage, 179, 225–234. https://doi.org/10.1016/j.neuroimage.2018.06.050

Brang, D., Dai, Z., Zheng, W., & Towle, V. L. (2016). Registering imaged ECoG electrodes to human cortex: A geometry-based technique. Journal of Neuroscience Methods, 273, 64–73. https://doi.org/10.1016/j.jneumeth.2016.08.007

Cardinale, F., Cossu, M., Castana, L., Casaceli, G., Schiariti, M. P., Miserocchi, A., Fuschillo, D., Moscato, A., Caborni, C., Arnulfo, G., & Lo Russo, G. (2013). Stereoelectroencephalography: Surgical methodology, safety, and stereotactic application accuracy in 500 procedures. Neurosurgery, 72(3), 353–366. https://doi.org/10.1227/NEU.0b013e31827d1161

Chang, E. F. (2015). Towards Large-Scale, Human-Based, Mesoscopic Neurotechnologies. Neuron, 86(1), 68–78. https://doi.org/10.1016/j.neuron.2015.03.037

Dale, A. M., Fischl, B., & Sereno, M. I. (1999). Cortical surface-based analysis. I. Segmentation and surface reconstruction. NeuroImage, 9(2), 179–194. https://doi.org/10.1006/nimg.1998.0395

Destrieux, C., Fischl, B., Dale, A., & Halgren, E. (2010). Automatic parcellation of human cortical gyri and sulci using standard anatomical nomenclature. NeuroImage, 53(1), 1–15. https://doi.org/10.1016/j.neuroimage.2010.06.010

Dykstra, A. R., Chan, A. M., Quinn, B. T., Zepeda, R., Keller, C. J., Cormier, J., Madsen, J. R., Eskandar, E. N., & Cash, S. S. (2012). Individualized localization and cortical surface-based registration of intracranial electrodes. NeuroImage, 59(4), 3563–3570. https://doi.org/10.1016/j.neuroimage.2011.11.046

Elias, W. J., Fu, K.-M., & Frysinger, R. C. (2007). Cortical and subcortical brain shift during stereotactic procedures. Journal of Neurosurgery, 107(5), 983–988. https://doi.org/10.3171/jns.2007.107.5.983

Erhardt, J. B., Lottner, T., Pasluosta, C. F., Gessner, I., Mathur, S., Schuettler, M., Bock, M., & Stieglitz, T. (2020). Fabrication and validation of reference structures for the localization of subdural standard- and micro-electrodes in MRI. Journal of Neural Engineering, 17(4), 046044. https://doi.org/10.1088/1741-2552/abad7a

Fang, Q., & Boas, D. A. (2009). Tetrahedral mesh generation from volumetric binary and grayscale images. In 2009 IEEE International Symposium on Biomedical Imaging: From Nano to Macro (pp. 1142–1145). IEEE.

Frauscher, B., von Ellenrieder, N., Zelmann, R., Doležalová, I., Minotti, L., Olivier, A., Hall, J., Hoffmann, D., Nguyen, D. K., Kahane, P., Dubeau, F., & Gotman, J. (2018). Atlas of the normal intracranial electroencephalogram: neurophysiological awake activity in different cortical areas. Brain, 141(March), 1–15. https://doi.org/10.1093/brain/awy035

Gupta, D., Hill, N. J., Adamo, M. a., Ritaccio, A., & Schalk, G. (2014). Localizing ECoG electrodes on the cortical anatomy without post-implantation imaging. NeuroImage: Clinical, 6, 64–76. https://doi.org/10.1016/j.nicl.2014.07.015

Hamilton, L. S., Chang, D. L., Lee, M. B., & Chang, E. F. (2017). Semi-automated Anatomical Labeling and Inter-subject Warping of High-Density Intracranial Recording Electrodes in Electrocorticography. Frontiers in Neuroinformatics, 11(October). https://doi.org/10.3389/fninf.2017.00062

Jiang, T., Liu, S., Pellizzer, G., Aydoseli, A., Karamursel, S., Sabanci, P. A., Sencer, A., Gurses, C., & Ince, N. F. (2018). Characterization of Hand Clenching in Human Sensorimotor Cortex Using High-, and Ultra-High Frequency Band Modulations of Electrocorticogram. Frontiers in Neuroscience, 12, 110. https://doi.org/10.3389/fnins.2018.00110

Katsura, M., Sato, J., Akahane, M., Kunimatsu, A., & Abe, O. (2018). Current and novel techniques for metal artifact reduction at CT: Practical guide for radiologists. Radiographics, 38(2), 450–461. https://doi.org/10.1148/rg.2018170102

Lachaux, J. P., Rudrauf, D., & Kahane, P. (2003). Intracranial EEG and human brain mapping. Journal of Physiology Paris, 97(4–6), 613–628. https://doi.org/10.1016/j.jphysparis.2004.01.018

LaPlante, R. A., Tang, W., Peled, N., Vallejo, D. I., Borzello, M., Dougherty, D. D., Eskandar, E. N., Widge, A. S., Cash, S. S., & Stufflebeam, S. M. (2016). The interactive electrode localization utility: software for automatic sorting and labeling of intracranial subdural electrodes. International Journal of Computer Assisted Radiology and Surgery, 1–9. https://doi.org/10.1007/s11548-016-1504-2

LaViolette, P. S., Rand, S. D., Ellingson, B. M., Raghavan, M., Lew, S. M., Schmainda, K. M., & Mueller, W. (2011). 3D visualization of subdural electrode shift as measured at craniotomy reopening. Epilepsy Research, 94(1–2), 102–109. https://doi.org/10.1016/j.eplepsyres.2011.01.011

Li, G., Jiang, S., Chen, C., Brunner, P., Wu, Z., Schalk, G., Chen, L., & Zhang, D. (2020). IEEGview: An open-source multifunction GUI-based Matlab toolbox for localization and visualization of human intracranial electrodes. Journal of Neural Engineering, 17(1). https://doi.org/10.1088/1741-2552/ab51a5

Martin, S., Iturrate, I., Millán, J. del R., Knight, R. T., & Pasley, B. N. (2018). Decoding inner speech using electrocorticography: Progress and challenges toward a speech prosthesis. In Frontiers in Neuroscience (Vol. 12, Issue JUN, p. 422). Frontiers Media S.A. https://doi.org/10.3389/fnins.2018.00422

Narizzano, M., Arnulfo, G., Ricci, S., Toselli, B., Tisdall, M., Canessa, A., Fato, M. M., & Cardinale, F. (2017). SEEG assistant: a 3DSlicer extension to support epilepsy surgery. BMC Bioinformatics, 18(1), 124. https://doi.org/10.1186/s12859-017-1545-8

Parvizi, J., & Kastner, S. (2017). Human intracranial EEG: Promises and Limitations. Nature Neuroscience. https://doi.org/10.1038/s41593-018-0108-2

Qin, C., Tan, Z., Pan, Y., Li, Y., Wang, L., Ren, L., Zhou, W., & Wang, L. (2017). Automatic and Precise Localization and Cortical Labeling of Subdural and Depth Intracranial Electrodes. Frontiers in Neuroinformatics, 11(February), 1–10. https://doi.org/10.3389/fninf.2017.00010

Rosenow, F., & Lüders, H. (2001). Presurgical evaluation of epilepsy. Brain, 124(9), 1683–1700. https://doi.org/10.1093/brain/124.9.1683

Rusinkiewicz, S. (2004). Estimating curvatures and their derivatives on triangle meshes. Proceedings - 2nd International Symposium on 3D Data Processing, Visualization, and Transmission. 3DPVT 2004, 486–493. https://doi.org/10.1109/TDPVT.2004.1335277

Stead, M., Bower, M., Brinkmann, B. H., Lee, K., Marsh, W. R., Meyer, F. B., Litt, B., Van Gompel, J., & Worrell, G. a. (2010). Microseizures and the spatiotemporal scales of human partial epilepsy. Brain : A Journal of Neurology, 133(9), 2789–2797. https://doi.org/10.1093/brain/awq190

Stolk, A., Griffin, S., Van Der Meij, R., Dewar, C., Saez, I., Lin, J. J., Piantoni, G., Schoffelen, J. M., Knight, R. T., & Oostenveld, R. (2018). Integrated analysis of anatomical and electrophysiological human intracranial data. Nature Protocols, 13(7), 1699–1723. https://doi.org/10.1038/s41596-018-0009-6

Studholme, C., Hill, D. L. G., & Hawkes, D. J. (1999). An overlap invariant entropy measure of 3D medical image alignment. Pattern Recognition, 32(1), 71–86. https://doi.org/10.1016/S0031-3203(98)00091-0

Studholme, C., Novotny, E., Zubal, I. G., & Duncan, J. S. (2001). Estimating tissue deformation between functional images induced by intracranial electrode implantation using anatomical MRI. NeuroImage, 13(4), 561–576. https://doi.org/10.1006/nimg.2000.0692

Taimouri, V., Akhondi-Asl, A., Tomas-Fernandez, X., Peters, J. M., Prabhu, S. P., Poduri, A., Takeoka, M., Loddenkemper, T., Bergin, A. M. R., Harini, C., Madsen, J. R., & Warfield, S. K. (2014). Electrode localization for planning surgical resection of the epileptogenic zone in pediatric epilepsy. International Journal of Computer Assisted Radiology and Surgery, 9(1), 91–105. https://doi.org/10.1007/s11548-013-0915-6

Trotta, M. S., Cocjin, J., Whitehead, E., Damera, S., Wittig, J. H., Saad, Z. S., Inati, S. K., & Zaghloul, K. A. (2018). Surface based electrode localization and standardized regions of interest for intracranial EEG. Human Brain Mapping, 39(2), 709–721. https://doi.org/10.1002/hbm.23876

Vakharia, V. N., Sparks, R., O’Keeffe, A. G., Rodionov, R., Miserocchi, A., McEvoy, A., Ourselin, S., & Duncan, J. (2017). Accuracy of intracranial electrode placement for stereoencephalography: A systematic review and meta-analysis. Epilepsia, 1–12. https://doi.org/10.1111/epi.13713

Wittek, A., Miller, K., Kikinis, R., & Warfield, S. K. (2007). Patient-specific model of brain deformation: Application to medical image registration. Journal of Biomechanics, 40(4), 919–929. https://doi.org/10.1016/j.jbiomech.2006.02.021

